# Species traits shape the intercontinental introduction patterns of bark and ambrosia beetles

**DOI:** 10.64898/2026.07.20.738676

**Authors:** Gimena Vilardo, Juan C. Corley, Andrew M. Liebhold, Eckehard G. Brockerhoff, Rebecca M. Turner, Takehiko Yamanaka, Sergio A. Estay, M. Victoria Lantschner

**Affiliations:** Grupo de Ecología de Poblaciones de Insectos, Instituto de Investigaciones Forestales y Agropecuarias Bariloche, INTA Bariloche-CONICET, Bariloche, Argentina; Faculty of Forestry and Wood Sciences, Czech University of Life Sciences, 165 00 Prague, Czech Republic; Swiss Federal Institute for Forest, Snow and Landscape Research WSL, 8903 Birmensdorf, Switzerland; Bioeconomy Science Institute, Tuhiraki, 19 Ellesmere Junction Road, Lincoln, 7608, New Zealand; Research Center for Agricultural Information Technology, National Agriculture and Food Research Organization, Tsukuba 305-0856, Japan; Universidad Austral de Chile, Instituto de Ciencias Ambientales y Evolutivas, Valdivia, Chile; Departamento de Ecología, CRUB Universidad Nacional del Comahue, Bariloche, Argentina

**Keywords:** Alien species, biotic homogenization, forests insects, global trade, invasions, propagule pressure, transport

## Abstract

1. A negative consequence of international trade is the accidental transport of non-native insect species to non-native regions, but our understanding of how this risk varies among species and how it relates to species traits remains incomplete.
2. We conducted a multi-continental analysis of historical border interception records for bark and ambrosia beetles (Coleoptera: Curculionidae: Scolytinae and Platypodinae), integrating 13,124 interception events (353 species) from Canada, Chile, Japan, New Zealand, and the United States. We estimated an intercontinental propagule pressure index and tested how species characteristics influence transport likelihood.
3. Interception assemblages showed low similarity among countries, with 64% of species recorded in only one nation, indicating strong geographic structuring linked to trade pathways. Interceptions were taxonomically and biogeographically non-random: species native to the Palearctic Nearctic and Indo-Malayan regions were overrepresented, whereas Neotropical, Austro-Pacific and Afrotropical taxa were underrepresented.
4. Host tree taxa emerged as the strongest predictor of propagule pressure. Species associated with conifers—particularly Pinaceae—exhibited approximately twice the propagule pressure of species restricted to broadleaved hosts. Ambrosia beetles showed higher propagule pressure than bark and twig/seed feeders. Propagule pressure also increased with the number of regions where species have already invaded, consistent with bridgehead dynamics.
5. Synthesis and applications: Our findings demonstrate that intercontinental transport of bark and ambrosia beetles is strongly structured by host associations, feeding guild, and biogeographic origin, reflecting the composition of traded commodities and historical trade networks. Identifying taxa with consistently high propagule pressure provides a basis for prioritizing surveillance and strengthening global forest biosecurity.

## Introduction

Invasions by exotic species remain one of the greatest environmental threats due to their high economic, social, and environmental impacts (Pyšek et al., 2020; Diagne et al., 2021). With increasing global trade and human travel, hundreds of species are unintentionally transported and can arrive in distant areas that they would otherwise be unable to reach on their own (Essl et al., 2015; Seebens et al., 2025). Forest insects are among organisms most frequently moved through international trade, as many species can be readily transported with imported live plants and with wood and wood packaging materials used in international commerce (Marini et al., 2011; Meurisse et al., 2018).

To prevent the introduction and spread of non-native, potentially invasive species, most countries implement a wide range of measures, including phytosanitary treatments of internationally traded materials, import prohibitions, border inspections, and traps at international entry points, among others (Nahrung et al., 2023; Ormsby and Brenton-Rule, 2017). By limiting the transport and arrival of potentially invasive organisms, these measures reduce propagule pressure (i.e., arrival rate), one of the strongest drivers of invasion success. Because eradication effort and costs increase as invasion progresses, preventing introductions is widely regarded as one of the most effective strategies for managing biological invasions (Leung et al., 2002; Epanchin-Niell et al., 2017).

Propagule pressure is the strongest driver of invasion success at the global scale (Brockerhoff and Liebhold, 2017; Nahrung et al., 2023; Vilardo et al., 2022). Propagule pressure of forest insects is determined by several factors, including the nature and volume of traded materials that act as vectors, the intensity of international trade, the effectiveness of phytosanitary measures, and species traits (Renault et al., 2018, Nahrung & Carnegie, 2021; Haack et al., 2014; Seebens et al., 2018). These drivers operate jointly to determine both the frequency and diversity of species moved across regions. Among these factors, species-specific biological and ecological characteristics play a key role in mediating different invasion stages (Chown and McGeoch 2023).

The probability of unintentional transport in wood packaging materials used in global trade usually increases for species with small body size and cryptic habits that allow them to remain unnoticed (e.g., under the bark or inside wood), broad host ranges, tolerance to desiccation or low temperatures, the ability to survive in dry or processed wood, long-lived or dormant life stages, and their association with traded commodities (Haack 2006; Renault et al., 2018; Meurisse et al., 2019; Grégoire et al., 2020; Nahrung and Carnegie 2021; Vilardo et al., 2022). However, the relative importance of these traits likely varies among taxonomic groups within forest insects. A deeper understanding of how transport-related characteristics operate across specific taxa is therefore essential to identify those insects most likely to be moved through global trade and consequently improving prevention strategies (Brockerhoff and Liebhold 2017; Grousset et al., 2020; Nahrung and Carnegie, 2021).

Bark and ambrosia beetles (Coleoptera: Curculionidae: Scolytinae and Platypodinae) constitute a large and diverse group of forest insects that exhibit a wide range of habits and exploit various plant tissues. Several species can cause severe forest impacts due to their ability to mass-attack and kill host trees (Biedermann et al., 2019; Raffa et al., 2008). These insects are commonly transported in raw logs and wood packaging materials (Brockerhoff et al., 2006; Meurisse et al., 2019), and consequently, numerous species have been intercepted at borders, several of which have established beyond their native ranges, in some cases becoming significant pests (Brockerhoff et al., 2014; Haack and Rabaglia, 2013; Faccoli et al., 2020; Lantschner et al., 2020; Gregoire et al., 2023). However, there is substantial variability in how often these species are introduced outside their native range (Brockerhoff et al., 2003, 2006, 2014; Haack 2001, 2006). This variability makes bark and ambrosia beetles an excellent model for exploring which traits enable some species to experience higher propagule pressure.

Certain traits of bark and ambrosia beetles may influence their likelihood of transport through international trade, leading to non-random interception patterns. Consequently, identifying these traits across species intercepted in different regions of the world is key to predicting their risk of introduction. Addressing this gap requires broader-scale studies that include countries from all continents. The frequency of interception of individual species during biosecurity inspections provides an indirect measure of the association with invasion pathways and serves as a proxy for their propagule pressure (Brockerhoff et al., 2014; Turner et al., 2021). The aims of this study are to describe the main patterns of interceptions of bark and ambrosia beetle species in imported products across five countries located in different continents, and to analyze how propagule pressure is influenced by the species characteristics.

## Materials and methods

### Interceptions databases

We obtained databases recording port interceptions of bark and ambrosia beetles (hereafter BAB; Coleoptera, Curculionidae, Scolytinae and Platypodinae) in imported products through maritime, air and terrestrial transport and their packaging material (pallets, containers, wooden crates) from five countries: Chile (1995-2005), Japan (1999-2017), New Zealand (1950-2008), the United States (1914-2008), and Canada (1997-2019) (see Supplementary Table S1 for more detail).

We compiled a list of the intercepted species in each country (hereafter “recipient countries”). Only interceptions identified to the species level were included in the analysis. To standardize taxonomic classifications across databases and periods covered, we used the most up-to-date scientific names for all recorded species, following Atkinson et al. (2026) and Alonzo-Zarazaga et al., 2023.

For each intercepted species in each recipient country, we estimated the interception frequency by summing all interception records over the entire period available for each recipient country. In most cases, the databases reported species interceptions without specifying the number of individuals detected. Therefore, an interception event was defined as the standard detection unit across all databases, regardless of the propagule size.

### Species characteristics

For each intercepted species, we collected information on its geographic distribution (native and non-native), feeding habit, host breadth and host taxa (Supplementary Table S2):

#### Geographic distribution

We determined the native biogeographic region of each species based on the countries where they are cited as native. For this purpose, we primarily consulted catalogs (Wood and Bright 1992; Knížek 2011; Alonso-Zarazaga et al., 2023) and online databases (e.g., Atkinson et al., 2026; Smith et al., 2019), supplemented this information with species-specific publications when necessary. We followed Wallace’s classification of biogeographic regions, with slight modifications to match the boundaries of the regions with the country’s borders. The regions considered were: Nearctic, Neotropical, Palearctic, Indo-Malayan, Afrotropical, and Austro-Pacific (see Lantschner et al., 2020 for more details). Species whose native biogeographic region could not be determined due to their widespread distribution and uncertain native range were classified as “cosmopolitan”. For species whose native distribution spanned more than one biogeographic region, we assigned the region where they are predominantly distributed to simplify the analysis.

For each species, we also collected information on whether it has established populations outside its native range, using available bibliographic sources (Wood & Bright 1992, Lantschner et al., 2020, Gregoire et al., 2023). We recorded the countries where each species is established and based on this information, we calculated the number of biogeographic regions in which each species occurs (including native and non-native ones). Using the same sources, we determined which species are currently established in each recipient country and calculated the percentage of those species that were intercepted during the study period.

#### Feeding habit

We classified species into three types of feeding habits: (1) bark beetles (BB), species that mainly feed on the bark or phloem (i.e., phloeophagy) of tree stems or branches; (2) ambrosia beetles (AB), species that primarily feed on fungi that they cultivate within the galleries they excavate in the xylem (i.e., xylomycetophagy); (3) other: including twig feeders, species feeding in pith of twigs, small branches or small stems (i.e. myelophagy), seed feeders, species feeding in large hard seeds and the encompassing fruit tissues (i.e., spermatophagy), and/ or herbivores, species that feed on other tissues of plants such as roots or leaves (i.e., herbiphagy) (Haack and Rabaglia, 2013; Kirkendall et al., 2015; Vega et al., 2015).

#### Host breadth and host taxa

We compiled information on the host plant families used by each species, to characterize their host taxa and breadth. We classified the species into three categories: those that use broadleaved trees, those that use conifers, and those that use both. This information was primarily obtained from Wood and Bright (1992), Atkinson (2024) and supplemented with specific publications for the species for which information was not available. Additionally, species were categorized based on their host breadth as “oligophagous”, if they used one or two plant families, or “polyphagous”, if they utilized more than two families.

### Data analysis

#### Patterns in intercepted species traits

We calculated the Cao index to quantify similarity between each pair of recipient countries in terms of intercepted species. The Cao index has been suggested as a minimally biased measure for cases of high beta diversity and variable sampling intensity (Cao et al., 1997).

For each species trait (geographic distribution, feeding habit, host breadth and host taxa) we analyzed whether the proportion of intercepted species within each category differed from their proportion observed among all native species using a Chi-squared test. We obtained information on known species diversity by biogeographic region from Hulcr et al. (2015) and on global diversity of feeding habits from Kirkendall et al. (2015). Information on the global diversity of species within each host breadth category and host taxa was compiled host catalogs (Wood and Bright 1992; Marchioro et al., 2024a, 2024b; Marchioro et al., 2025; Ruzzier et al., 2023; Šenfeldová et al., 2024) and online databases (e.g., Atkinson et al., 2026).

We also tested whether proportions of intercepted species of each trait category varied significantly among recipient countries using a chi-square test. Since the chi-squared test requires that expected frequencies are sufficiently large for valid approximation (Agresti, 2002), categories that did not fit this requirement were excluded from the analysis.

#### Drivers of intercontinental propagule pressure

We estimated an index of “intercontinental propagule pressure” of BAB species, that integrates both the interception frequency within each country and the frequency with which each species is intercepted across all countries. This index was calculated as the Shannon index for each species, considering the number of interception events in each country and the number of countries where the species was intercepted. We used the following: 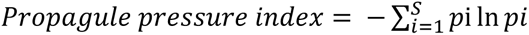, where S is the number of recipient countries, *p*i is the proportion of interception events of species *i,* relative to the total number of interception events in each recipient.

We fitted a generalized linear model (GLM) to analyze how the propagule pressure index varied among the intercepted species based on their traits: Native biogeographic region, number of regions where the species occur (including native and non-native ones), feeding habit, host breadth and host taxa. Prior to analysis, the index was transformed using a lognormal distribution as it was determined to be the most suitable after evaluating alternative continuous variables distributions using the “*fitdistrplus*” function (Delignette-Muller, 2015). We calculated the adjusted *R^2^* and the root mean squared error (RMSE) as a metric of goodness of fit.

## Results

A total of 353 bark and ambrosia beetle species were recorded from a total of 13,124 interceptions among five recipient countries. This represents approximately 5% of the 7,400 species described globally for the Platypodinae and Scolytinae subfamilies (Kirkendall et al., 2015). Eighty-five percent of the intercepted species belonged to the subfamily Scolytinae, and 15% to the subfamily Platypodinae. Japan was the country that intercepted the most species (187) followed by the USA (172), New Zealand (112), Canada (82), and Chile (44). Across the five countries analyzed, a substantial proportion of bark and ambrosia beetle interceptions were identified only at the family level, ranging from 25% in Japan to 79% in Chile, with intermediate values in Canada (29%), New Zealand (26%), and the USA (55%); in Canada, an additional 23% of intercepted Curculionidae were identified only to family level, which implies that the real number of intercepted species is much larger.

### Similarity in intercepted species among countries

Among the intercepted species, 64% (227) were recorded in only one country. A total of 18 species (5%) were intercepted in all five recipient countries (Table S2). The similarity matrix based on Cao’s index revealed relatively low similarity between countries in the composition of intercepted species, with values ranging from 0 to 0.19. The USA and New Zealand showed the highest similarity among countries, while Japan and Chile exhibited no overlap (Figure 1).

**Figure 1.**
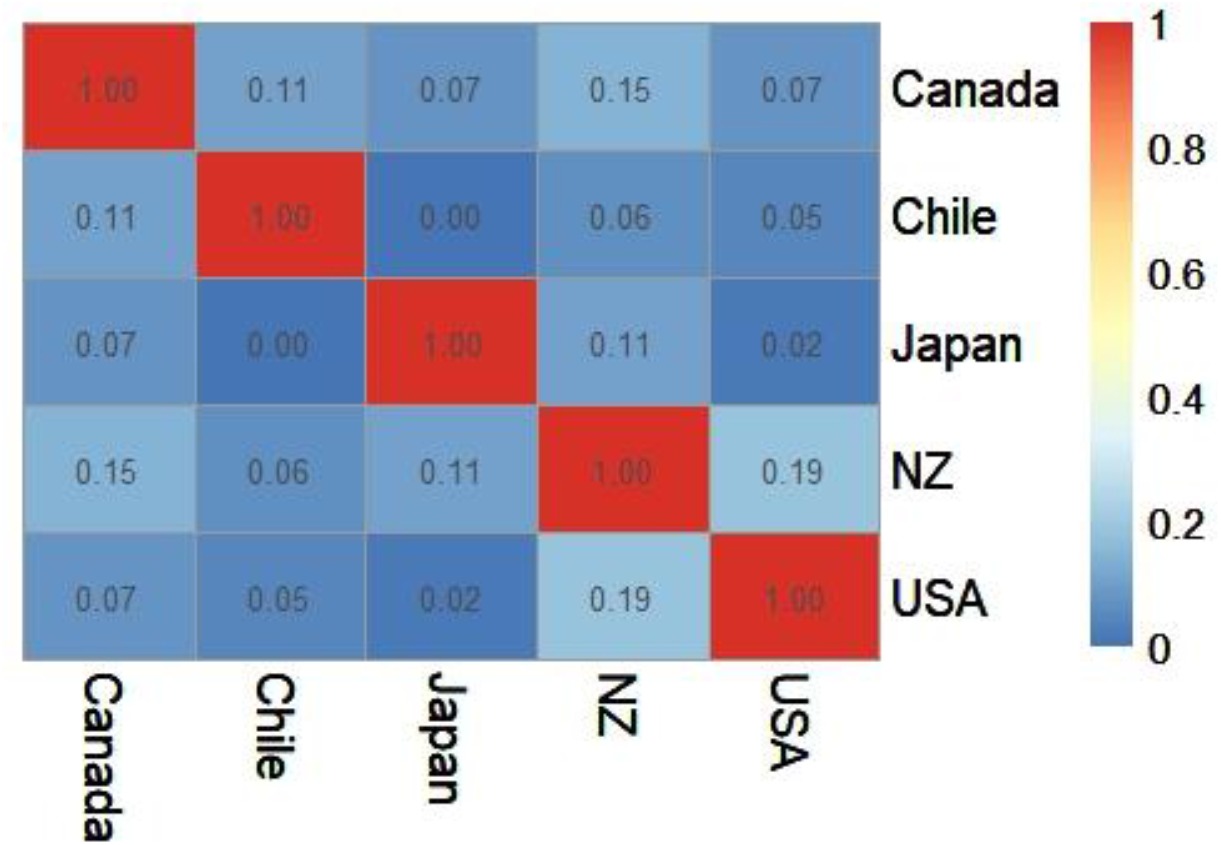
Heatmap showing the similarity in the species composition of intercepted bark and ambrosia beetle assemblages among Canada, Chile, Japan, New Zealand and the United States, based on Cao’s index. Values range from 0 (no similarity) to 1 (red indicating complete similarity in the diagonal, when compared with their own).

### Patterns in intercepted species traits

#### Geographic distribution

Most intercepted species were native to the Palearctic region (34%), followed by the Nearctic (24%) and the Indo-Malayan (24%) regions (Figure 2A). These proportions differed significantly from those based on frequencies of native species in each region (*X^2^*: inf, p <0.01). The Palearctic, Nearctic and Indo-Malayan regions were overrepresented among interceptions, whereas the Neotropical, Afrotropical, and Austro-Pacific regions were underrepresented (Supplementary Table S3).

**Figure 2.**
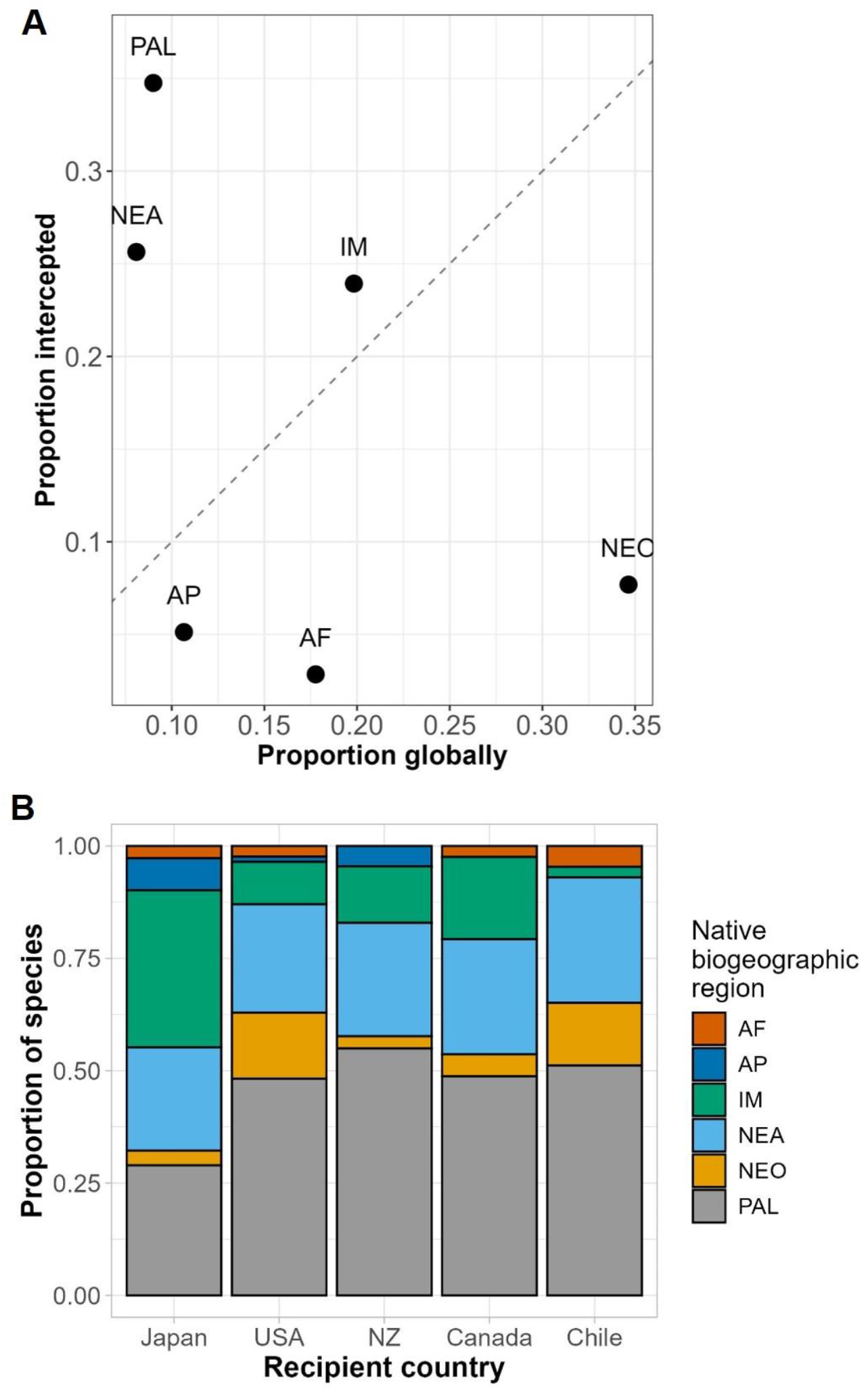
Native biogeographic region of the intercepted species. (A) Proportion of intercepted species and known species in native to each biogeographic region. (B) Proportion of intercepted species from each native biogeographic region in each recipient country. AF: Afrotropical, AP: Austro-Pacific, NEO: Neotropic, IM: Indo-Malayan, NEA: Nearctic, PAL: Palearctic.

Proportions of intercepted native species from each biogeographic region varied among the five recipient countries (*X^2^: 98*, p < 0.001). In Chile, Canada, New Zealand, and the USA, interceptions were dominated by species native to the Palearctic region, followed by those from the Nearctic region. In contrast, interceptions in Japan were dominated by species native to the Indo-Malayan region, although species native to the Palearctic and Nearctic regions were also important. In all countries, interceptions of species native to the Afrotropical and Austro-Pacific regions were very low, whereas those of Neotropical origin were more variable, being more important in the USA and Chile. Canada and Chile did not intercept species native to the Austro-Pacific region, nor did New Zealand intercept species native from the Afrotropical region (Figure 2B).

Most intercepted species were not native to the recipient country (between 70% and 91%). Among the non-native species intercepted, only a small fraction were established in the recipient country (ranging from 2% in Japan, 4% in NZ, 16% in Canada and Chile and 22% in USA) (Figure 3). The proportion of species currently reported as established in each recipient country that were intercepted during the study period varied among countries: 25% (2 of 8 species) in Japan, 32% (24 of 75 species) in the USA, 42% (5 of 12 species) in New Zealand, 44% (8 of 18 species) in Canada, and 46 % (6 of 13 species) in Chile (Supplementary Table S4).

**Figure 3.**
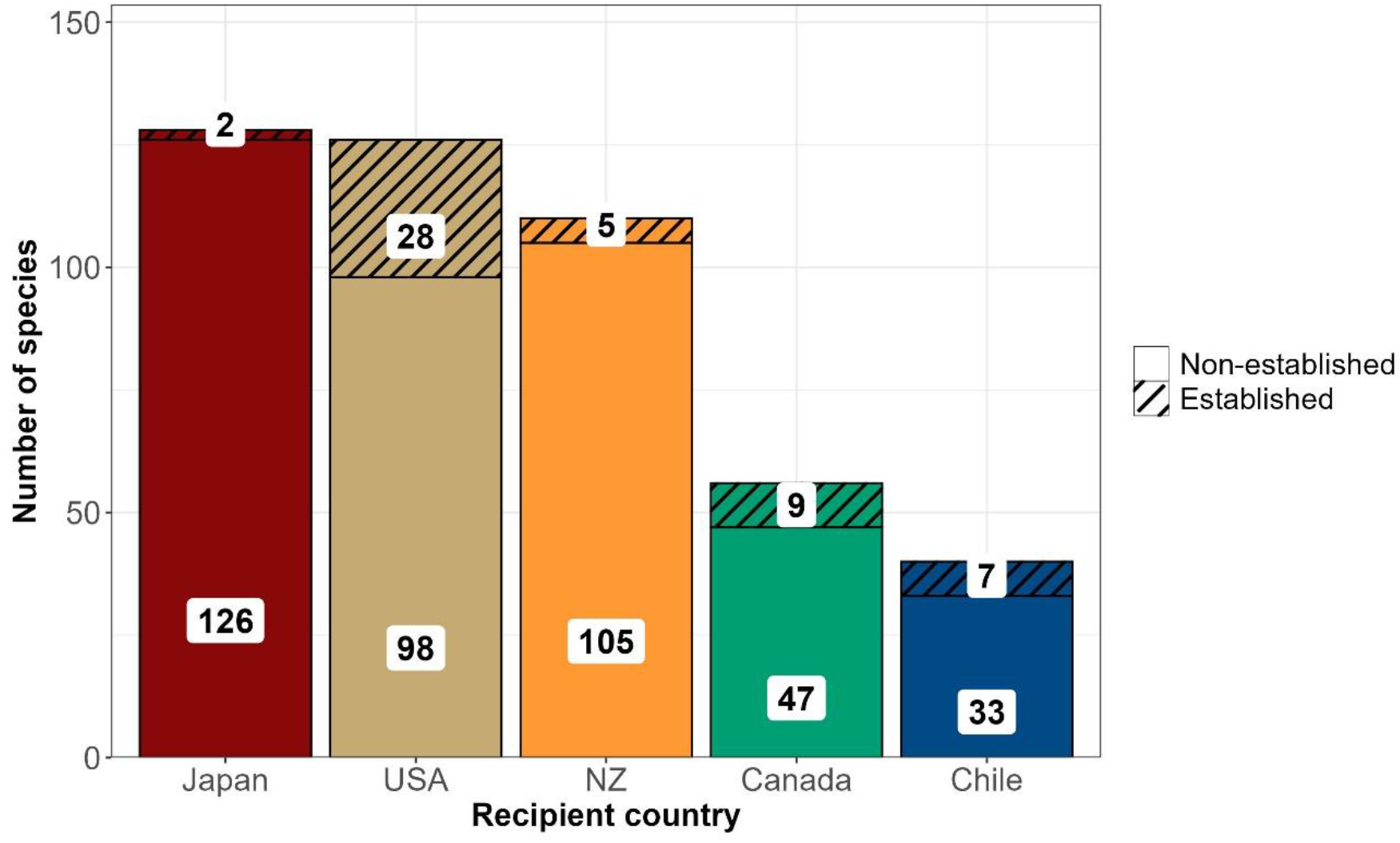
Number of intercepted non-native species intercepted in each recipient country. We indicate the number of species that have established in the recipient country versus those that have not.

#### Feeding habit

Fifty percent of the intercepted species were bark beetles, 41% were ambrosia beetles and 9% were seed and twig feeders. We found that the proportion of intercepted species in each feeding habit was very similar to proportions observed among species worldwide (Figure 4A; Supplementary Table S3) (*X^2^: 2*, p = 0.3). However, proportions in each category among countries (*X^2^: 43*, p < 0.001). In Japan, ambrosia beetles dominated; in the USA and New Zealand, bark beetles dominated, while in Chile and Canada, proportions of ambrosia beetles and bark beetles were similar. In all cases, the “other” feeding type represented a very low proportion (Figure 4B).

**Figure 4.**
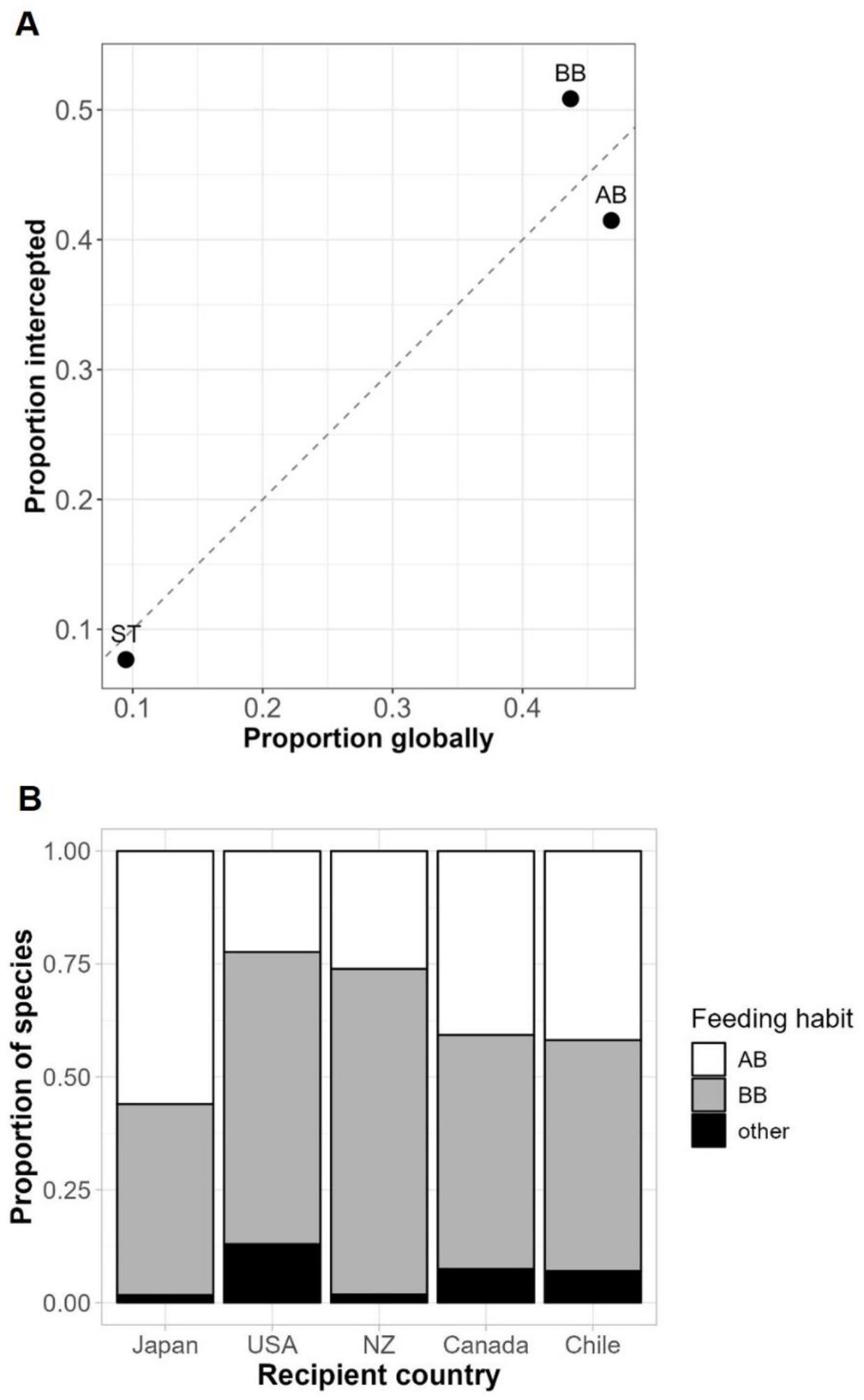
Feeding habit of intercepted species. (A) Proportions of intercepted species and native species belonging to each feeding habit. (B) Proportion of the total number of intercepted species by feeding habits in each recipient country. AB: ambrosia beetles; BB: bark beetles; other (twig, seed, and/ or leaf feeders). *Sauroptilius sauropterus* could not be classified due to insufficient information.

#### Host breadth

Of all the intercepted species, 60% (212 species) were oligophagous, 35% (122 species) were polyphagous, and 5% (19 species) had unknown hosts preferences. We found that the proportions of intercepted species in each host breadth category differed from proportions among all species globally (Figure 5A, Supplementary Table S3) (*X^2^: 50*, p = <0.001). Polyphagous species were overrepresented among intercepted species, and oligophagous species were underrepresented. The proportions of intercepted species across host breadth categories were similar among countries (*X^2^: 5.3*, p = 0.26; Figure 5B).

**Figure 5.**
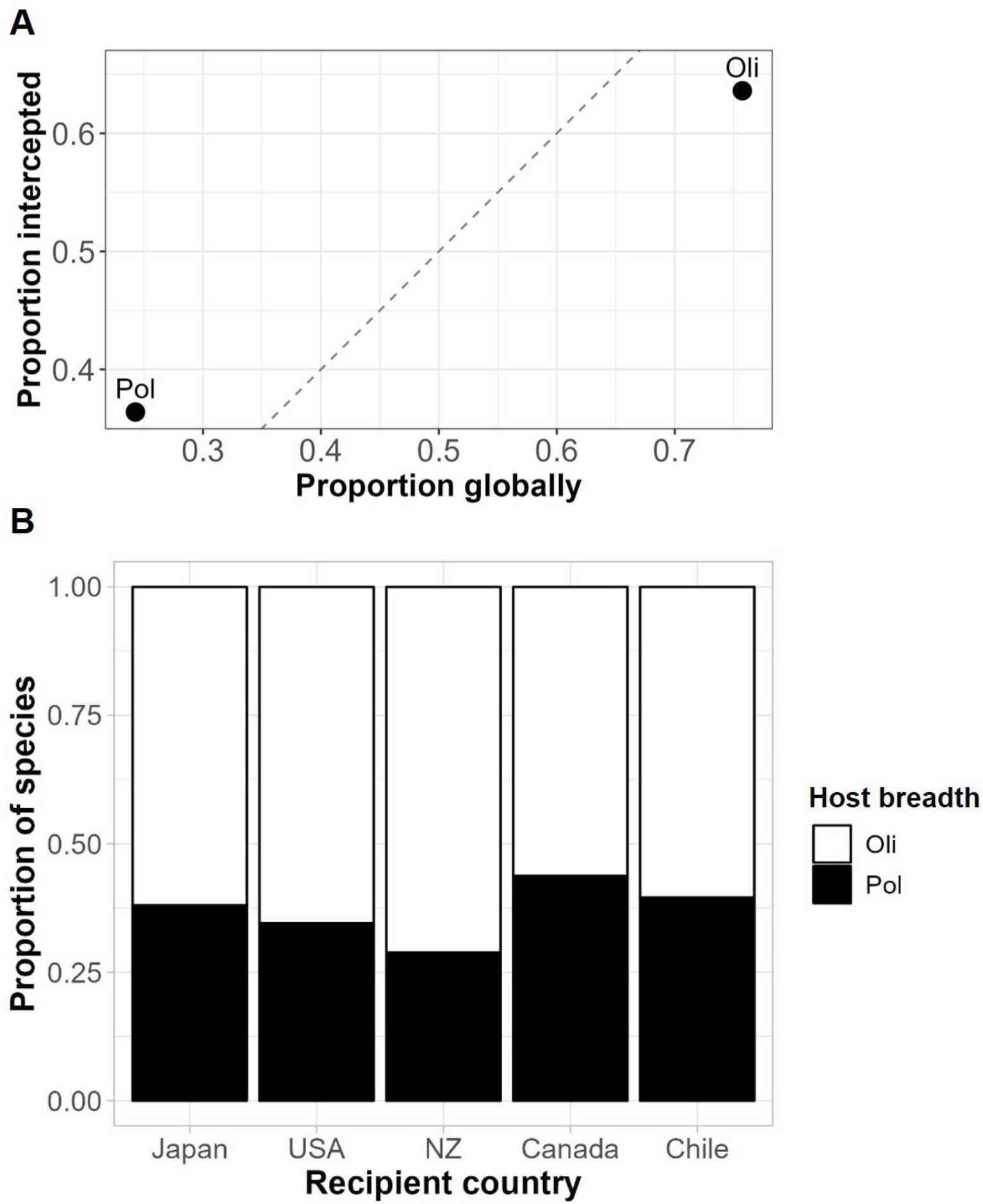
Host breadth among species. (A) Proportions of intercepted species and global species belonging to each host breadth category. Eighteen intercepted species could not be classified due to insufficient information. (B) Proportion of species in each country belonging to each host breadth category. Oli: oligophagous (one or two host plant families); Pol: polyphagous (more than two host families). Nineteen intercepted species could not be classified due to insufficient information.

#### Host taxa

Host groups varied according to the species’ host breadth. Among the oligophagous species, the most frequently utilized host family was Pinaceae (60% of the species), followed by Dipterocarpaceae (9%), Cupressaceae (6%), Moraceae (3%), and Oleaceae (3%). In turn, polyphagous species utilized 139 host families, with the most frequently used family being Fagaceae (51% of the species), Moraceae (41%), Dipterocarpaceae (37%), Lauraceae (37%), Anacardiaceae (34%) and Malvaceae (32.5%) (Supplementary Table S2).

When we classified species according to whether they utilize conifer hosts, broadleaf hosts, or both conifers and broadleaves, we found that the proportion of intercepted species in each category differed from the proportions observed among all world species (*X^2^:* 40, p <0.001). Species that use conifers, were significantly overrepresented among intercepted species, whereas species that use only broadleaved trees were underrepresented (Figure 6A, Supplementary Table S3).

**Figure 6.**
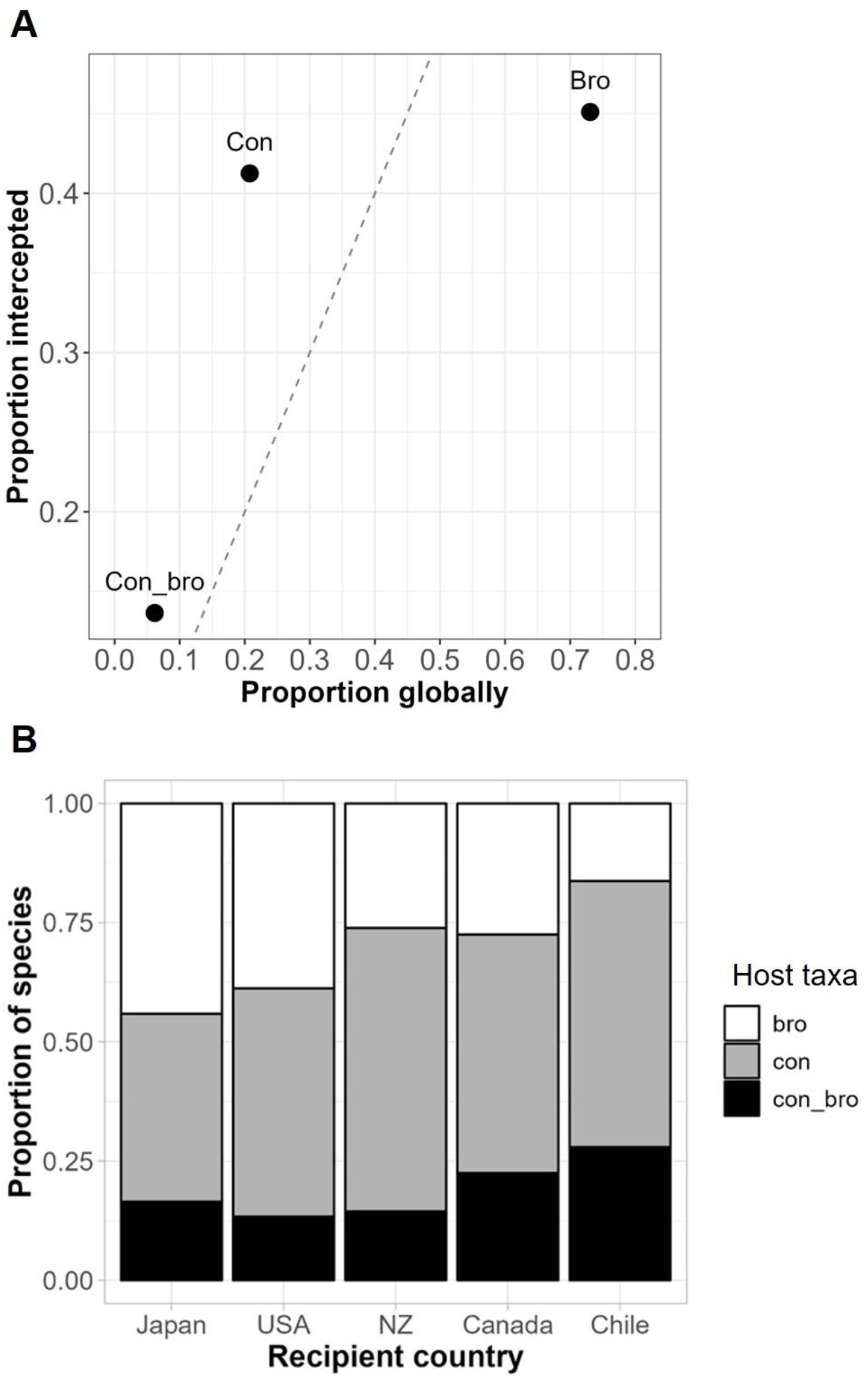
Host category of intercepted species. (A) Proportion of intercepted species and word species belonging to each host taxa category. (B) Proportion species in each country belonging to each host taxa category. Bro: use broadleaved trees as host; Con: use conifers, Con_bro: use both as hosts. Nineteen intercepted species could not be classified due to insufficient information.

Across recipient countries, the composition of intercepted species differed in the relative contributions of the three host-use categories (*X^2^: 25*, p <0.01; Figure 10). In Japan, the USA, New Zealand, and Canada, species associated exclusively with conifer hosts dominated interceptions, representing more than half of the species in each country. Species that utilize both conifers and broadleaf host species consistently accounted for a smaller but notable proportion (approximately 10–20%), whereas species restricted to broadleaf hosts represented the smallest share of interceptions (Figure 6B).

### Drivers of intercontinental propagule pressure

The propagule pressure index ranged from 0.001 to 0.75 among intercepted species. The first quartile of the distribution across species was 0.004, the median was 0.01, and 75% of the species had a propagule pressure index below 0.04. Species in the highest quartile of propagule pressure (0.04 to 0.075) included *Trypodendron lineatum*, followed by *Xyleborus perforans, Ips typographus, Pityogenes chalcographus, Hylurgops palliatus*, *Hylurgus ligniperda*, *Xyleborinus saxesenii and Orthotomicus erosus,* representing 2% of the species. The 25 species with a propagule pressure index higher than 0.15 accounted for 72% of the total interceptions. Eight of these species (30%) have not established populations outside their native range anywhere in the world, despite being frequently intercepted (Table S2).

Propagule pressure increased significantly with the number of regions in which a species was established, increasing by a factor of 1.62 for each additional region. Species associated exclusively with conifers exhibited propagule pressures 2.15 times higher than species associated exclusively with broadleaved hosts. Feeding habit also affected propagule pressure, with ambrosia beetles exhibiting higher propagule pressures than both bark beetles (1.68-fold higher) and seed- and twig-feeding beetles (4.32-fold higher). Native range had a relatively small effect, with only Neotropical species showing significantly lower propagule pressures than Afrotropical species. Host breadth did not have a significant effect explaining the propagule pressure. The model explained 23% of the variation in log-transformed propagule pressure (adjusted R² = 0.23; RMSE = 1.36), improving prediction accuracy relative to the null model (RMSE = 1.58) (Table 1, Figure 7).

**Figure 7.**
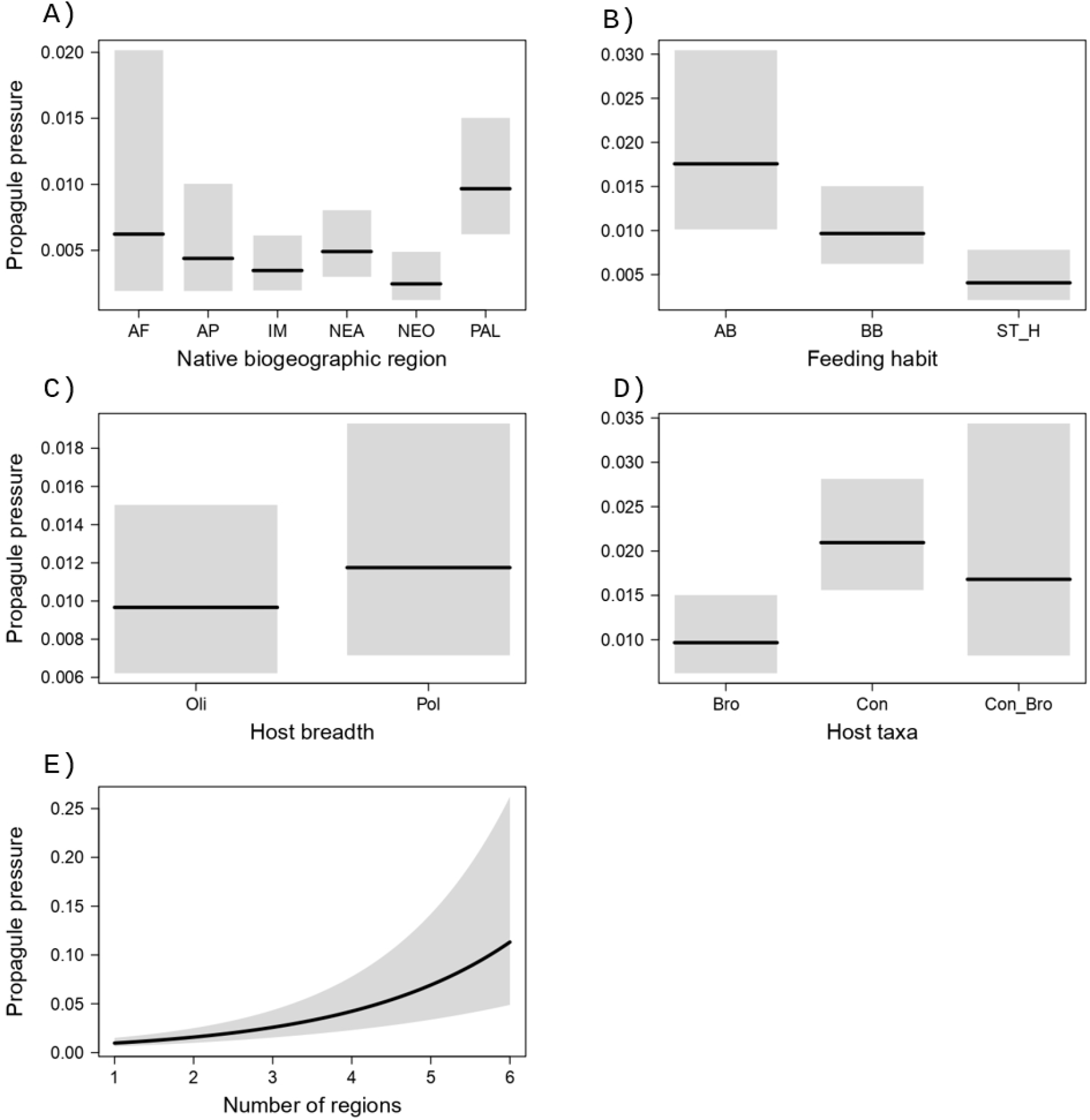
Mean propagule pressure index and 95% confidence intervals predicted by the model, considering partial effects of each predictor variable: (A) Species native range: AF (Afrotropic), AP (Austro-Pacific), IM (Indo-Malayan), NEA (Nearctic), NEO (Neotropic), PAL (Palearctic). (B) feeding habit: AB (ambrosia beetles), BB (bark beetles), ST_H (Seed, shoot, leaf and/ or root feeders). (C) Host breadth: Oli (oligophagous), Pol (polyphagous). (D) Host taxa: Bro (broadleaves), Con (conifers) and Bro_Con (broadleaves and conifers). (E) Number of regions where each species occurs (including native and non-native range).

**Table 1.**
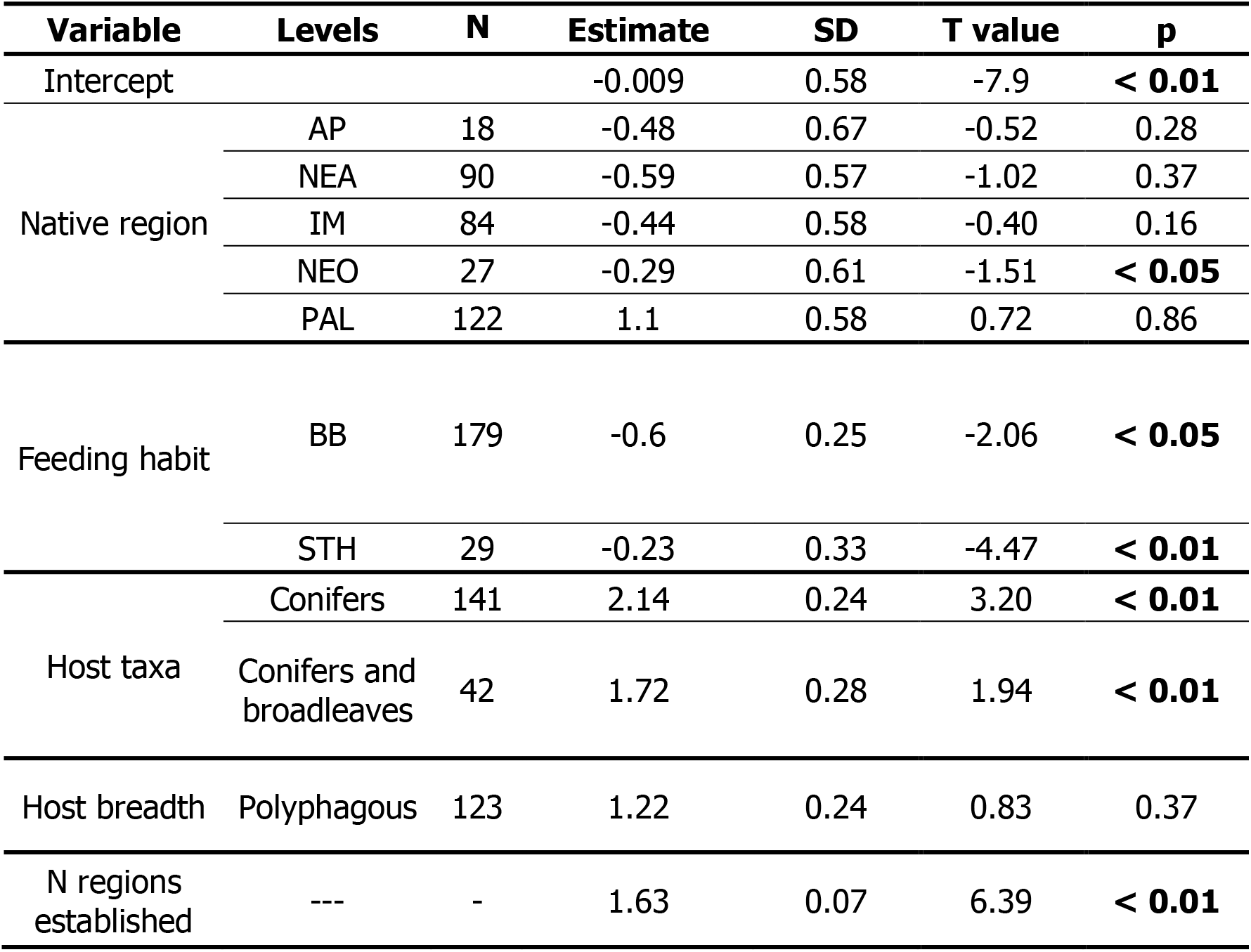
Effect size of the explanatory variables on the propagule pressure of the species. Explanatory variables were: feeding habit: AB (ambrosia beetles), BB (bark beetles), STB (Seeds and twig beetles); host taxa: broadleaves, conifers, and conifers and broadleaves; host breadth: Oli (oligophagous), Poli (polyphagous); number of regions where species are established; native range: AF (Afrotropic), AP (Austro-Pacific), NEA (Nearctic), IM (Indo-Malaya), NEO (Neotropical), PAL (Palearctic). Values are in the response scale (exp^(estimate)^). The intercept corresponds to AF, AB, Broadleaves and Oli species.

## Discussion

This study provides the first analysis of multi-continental variation in propagule pressure (proxied by border interceptions) of bark and ambrosia beetles. By integrating interception data from five countries spanning distinct biogeographic regions and comparing traits of intercepted species with their relative representation in native species pools, we confirm that propagule pressure is far from randomly distributed across taxa (Turner et al., 2021). The proportions of species traits observed among intercepted species differ markedly from those of all native species in the world, indicating that intercontinental transport is strongly filtered by species’ host associations, feeding habits, and geographical distribution, yielding consistent and predictable patterns in the likelihood of transport through global trade pathways.

Our results further suggest that introduction patterns of BAB generally differ from patterns observed in global native assemblages, such that larger native species pools do not necessarily translate into higher numbers of invasive species (Issit et al., 2024; Mally et al., 2021). This discrepancy aligns with the concept of invasion disharmony - a mismatch between native and exotic ensembles (Liebhold et al., 2021).

### Similarity in intercepted species among countries

Despite the highly globalized nature of trade, the species overlap among interception assemblages was remarkably low, even considering only a subset of the intercepted species. Approximately two-thirds of species were intercepted in only a single country, whereas only 5% were detected across all five. This pronounced geographic structuring indicates that interception datasets retain strong national signatures, likely reflecting differences in trade partners, commodity flows, and inspection histories, and suggests—consistent with previous studies— that introduced assemblages are shaped primarily by trade directionality and pathway specificity rather than by the global species pool (Lantschner et al., 2020; Liebhold et al., 2021; Fenn-Moltu et al., 2023; article in prep.). This trend contrasts with that observed in other taxonomic groups, such as Hemiptera and Thysanoptera, whose species interception frequencies were more similar across different regions of the world (Turner et al., 2021). However, these differences may be partly reflect variation in inspection priorities among taxonomic groups as several Hemiptera and Thysanoptera include well-known pest species that are specifically targeted during inspections, whereas bark and ambrosia beetles are not always priority targets in every country’s inspection programs. Comparisons among taxa under similar inspection efforts would improve the robustness of assessments such as ours and strengthen the conclusions that can be drawn from them (Brockerhoff et al., 2014). In addition, the low overlap observed here may be accentuated by incomplete sampling of arrival events, since interception records represent only a small fraction of actual introductions and are dominated by rarely intercepted species. Finally, taxonomic uncertainty and differences in identification capacity among countries may further contribute to underestimating the true degree of species overlap among interception assemblages.

Consequently, no single country’s interception data can adequately capture global transport risk of BAB, highlighting the value of multi-country syntheses —such as the one presented here— for identifying species of global invasion relevance. Although only a small proportion of described bark and ambrosia beetle species were intercepted across the five surveyed countries, incorporating additional national datasets would likely reveal a much broader set of transported taxa. Moreover, interception records represent only a fraction of species arriving at national borders (Turner et al., 2021, 2025), and a substantial proportion of bark and ambrosia beetle interceptions (25–79%, depending on the country) could not be identified to species level due to immature life stages, specimen damage, or taxonomic constraints. Together, these factors strongly suggest that both the number of species reported here and the similarity among species assemblages are underestimated.

At the species level, propagule pressure was highly skewed, with just 25 species accounting for 72% of all interceptions and only 2% of species falling within the highest quartile of propagule pressure. This skewed interception pattern is consistent with interception of other insect taxa (Turner et al., 2021). Though there is a positive association between interception frequency and establishment (Brockerhoff et al., 2006, 2014), high interception frequency did not consistently translate into establishment success in our analysis. Across the countries assessed here, most intercepted species (70–91%) were non-native, although these proportions should be interpreted cautiously because native species are not consistently recorded across interception programs. Only a small fraction (2–22%) have successfully invaded, and eight species exhibiting exceptionally high propagule pressure have never established outside their native ranges. This discrepancy underscores the fact that transport alone does not guarantee establishment, which is likely constrained by other factors such as climatic mismatch, Allee effects, limited host availability, or other ecological filters encountered at the destination (Vilardo et al., 2022). This is the case for example, of *Ips typographus*, which despite its high propagule pressure in USA it has not established yet in this country, probably due to strong Allee effects (Ward et al., 2023).

### Native range

Interceptions were dominated by species native to the Palearctic and Nearctic regions, far exceeding expectations based on their natural species richness. In contrast, species from the Neotropical, Afrotropical, and Austro-Pacific regions were strongly underrepresented. The patterns found here likely arise from two interacting mechanisms. On the one hand, Holarctic regions have historically dominated the global movement of people and goods, resulting in a disproportionate share of contemporary trade flows associated with the development of the industrial, agricultural and forestry sectors (Hulme 2009; López et al., 2023). In addition, Pinaceae species, which are exclusively native to the Northern Hemisphere, are widely used in packaging materials (Krishnankutty et al., 2020), represent a major transport pathway for bark and woodboring insects (Meurisse et al., 2019). Because pine (and other conifers) is widely available for use in solid wood packaging material originating from both the Northern Hemisphere and the Southern Hemisphere, where it is extensively planted for industrial forestry (Payn et al., 2016), insect species using pine wood as a host can be expected to be ubiquitous in wood packaging material used in trade worldwide. This phenomenon likely explains both the predominance of insects using conifers as hosts and the dominance of insects native to the Northern Hemisphere.

Propagule pressure was positively associated with the number of biogeographic regions in which a species is already established, suggesting that it is likely that these species continue to disperse through trade (Brockerhoff et al., 2014; Grégoire et al., 2023, article in prep). Accordingly, regions colonized beyond a species’ native range may act as new sources of secondary invasions, a phenomenon known as the bridgehead effect (Lombaert et al., 2010; article in prep).

### Feeding habit

Overall, the representation of feeding guilds in terms of number of intercepted species broadly mirrored their natural diversity, although the known global diversity of bark and ambrosia beetles is likely underestimated, as hundreds of species are believed to remain undescribed, particularly in tropical regions (Haack and Rabaglia, 2013). However, clear differences emerged among countries, with ambrosia beetles dominating interceptions in Japan, whereas bark beetles were more frequently intercepted in the USA and New Zealand. In contrast to species richness, interception frequency (i.e., propagule pressure) differed markedly among feeding guilds, with ambrosia beetles exhibiting the highest values, exceeding those of both bark beetles and twig/seed feeders. This pattern might reflect differences in transport dynamics among feeding guilds. Ambrosia beetles might experience greater exposure to internationally traded live plants, wood and wood-packaging materials, reflecting their close association with processed wood substrates.

In addition, ambrosia beetle species intercepted in this study tended to exhibit broader global distributions than true bark beetles. Approximately 10% of the ambrosia beetle species occurred in three or more biogeographic regions, compared with only 4% of bark beetles. This pattern suggests that intercepted ambrosia beetles are, on average, more geographically widespread, which may further contribute to their repeated detection across multiple recipient countries through secondary introductions (article in prep.). A likely explanation is that many ambrosia beetles exhibit sib-mating or inbreeding mating systems, allowing a single fertilized female to establish a new population. As a result, these species can successfully invade even under very low propagule pressure, in contrast to most bark beetles, which generally require multiple individuals to found a viable population (Kirkendall, 1983).

### Host breadth and category

Although oligophagous species constituted the largest proportion of intercepted taxa, polyphagous species were overrepresented relative to their global frequency across countries. Additionally, species that use conifers were overrepresented among intercepted species relative to those that use broadleaf hosts. Previous studies have found that species with greater host breadth are generally more frequent among established non-native species (Hulcr et al., 2007; Šenfeldová et al., 2024) though these studies have not differentiated the effect of transport vs. establishment success. Nevertheless, host breadth *per se* did not influence propagule pressure of intercepted species. This apparent disagreement likely reflects the characteristics of international wood trade: although a wide diversity of tree species is used in the manufacture of wood products and packaging materials, a relatively small subset of tree genera is used disproportionately often (Krishnankutty et al., 2020).

It is important to note, however, that estimates of global host breadth and host-use patterns remain incomplete because host associations are still unknown for many bark and ambrosia beetle species (Supplementary Table S3). Most species lacking host information are ambrosia beetles, a group that is more frequently polyphagous and commonly associated with broadleaf hosts. Consequently, the global proportion of polyphagous species is likely underestimated, suggesting that the apparent overrepresentation of polyphagous species in interception records may be less pronounced than observed here. Conversely, because species lacking host records are less frequently associated exclusively with conifers, the overrepresentation of conifer-feeding species in interceptions relative to their global diversity may be even greater than currently estimated.

Consistent with this interpretation, host taxa emerged as the strongest predictor of propagule pressure among the species traits examined, with higher interception rates for conifer-feeding beetle species. This prominent role of host taxa indicates that interception patterns are driven less by intrinsic feeding specialization and more by the types of wood moving in international trade, such as the widespread use of conifer wood in packaging and pallets.

Pinaceae was by far the most frequently represented host family, particularly among oligophagous species, and species associated with conifers were disproportionately intercepted. This pattern is likely explained, in part, by the broad global distribution of Pinaceae, especially pines (*Pinus* spp.). Native Pinaceae forests are very common across the Holarctic region, supporting a high diversity of bark and ambrosia beetles associated with this family (Wood and Bright 1992). In addition, extensive plantations of non-native pines occur throughout the Southern Hemisphere (Procheş et al., 2012; FAO 2020), potentially facilitating the secondary spread of already established non-native populations (i.e., bridgehead effect, article in prep). This dynamic is further reinforced by the frequent use of Pinaceae timber in international wood packaging materials, increasing opportunities for transport and repeated introductions (Krishnankutty et al., 2020).

### Combined effects of species traits and implications for biosecurity

Our findings have implications for biosecurity and the design of surveillance systems. Interception patterns consistently highlight conifer-based wood packaging as a major high-risk pathway, particularly for shipments originating from Palearctic and Nearctic regions, emphasizing the need for strengthened and targeted inspections along these routes. Species exhibiting high propagule pressure emerge as priority targets for surveillance, especially those that are repeatedly intercepted yet remain absent from recipient regions (*Trypodendron lineatum, Dryocoetes autographus, Ips sexdentatus*), as they represent a substantial pool of future invaders.

The interception databases compiled from different countries span distinct time periods that are not fully comparable, and both sampling effort and inspection protocols vary substantially among jurisdictions (Brockerhoff et al., 2014). Consequently, sampling is not random and is often biased toward species and countries considered high-risk. In addition, interception of native species may not be consistently recorded. This is particularly relevant in Japan, where many Palearctic bark and ambrosia beetle species are also native, potentially contributing to the relatively low number of Palearctic species reported in interception records. Similar practices may occur in other countries, such as the USA, where native species are not routinely recorded. For this reason, variability in the observed patterns should be interpreted with caution.

It is also important to note that the historical interception records analyzed here were mostly collected prior to the widespread implementation of the ISPM-15 standards and reflect past trade configurations. As a result, subsequent changes in phytosanitary policies—such as the adoption of ISPM 15—and the emergence of new trade partnerships are likely to have modified the interception patterns identified here (Haack et al., 2014, 2022).

The high proportion of interceptions lacking species-level identification (25–79% across countries) denotes a major limitation, largely driven by taxonomic challenges, particularly for immature stages commonly transported in wood, as well as misidentification among morphologically similar adult species. This may bias the results, particularly by leading to the underrepresentation of poorly known and/or difficult-to-identify species; and consequently, can compromise risk assessment, leading to misdirected management efforts. Strengthening taxonomic capacity through updated training and resources, together with the broader implementation of novel identification tools such as molecular diagnostic, would substantially improve the speed and accuracy of identifications (Poland and Rassati 2019). Finally, the strong geographic structuring of interception data demonstrates that no single national database adequately captures global movement patterns, reinforcing the importance of enhanced international data sharing and coordinated monitoring efforts to effectively anticipate and mitigate emerging invasion risks.

Previous studies have examined how species traits influence the invasion success of BAB by jointly considering multiple invasion stages, including transport, establishment, and subsequent spread. This work has consistently shown that species traits play an important role in determining invasion outcomes, with larger body size and broader host ranges being associated with a higher likelihood of invasion success (Vilardo et al., 2022). Similarly, sib-mating or inbreeding, polyphagy, and associations with fungal symbionts have been identified as key traits facilitating invasion success in these groups (Grousset et al., 2020; Lantschner et al., 2020; Grégoire et al., 2023). Studies in other insect taxa have begun to separate the transport stage from subsequent invasion processes using interception data, revealing that the traits associated with successful transport may differ from those promoting establishment and spread (Mally et al., 2022; Fenn-Moltu et al., 2023; Liebhold et al., 2024). In contrast, our study provides the first such assessment for BAB species involved in intercontinental transport. We show that the intercontinental movement of bark and ambrosia beetles is shaped by predictable species traits, with conifer-associated species and ambrosia beetles contributing disproportionately to intercontinental propagule pressure, particularly those originating from Holarctic regions. However, these relationships likely reflect historical trade patterns and environmental conditions, and ongoing global change may alter the characteristics that enable species to overcome dispersal barriers (Caplat et al., 2016). By identifying groups of species that pose elevated introduction risk, our findings provide valuable information for improving global biosecurity strategies aimed at preventing future forest insect invasions.

## Author contributions

Gimena Vilardo, Victoria Lantschner and Juan C. Corley conceived the ideas and designed the methodology; Andrew M. Liebhold, Eckehard G. Brockerhoff, Rebecca M. Turner, Takehiko Yamanaka, Sergio A. Estay collected the data; Gimena Vilardo and Victoria Lantschner analysed the data; Gimena Vilardo and Victoria Lantschner led the writing of the manuscript. All authors contributed critically to the drafts and gave final approval for publication.

## Acknowledgements

GV, JC and MVL acknowledge support from the Agencia Nacional de Promoción Científica y Tecnológica (PICT 2019-235) and CONICET (PIP 11220200100764CO). AML acknowledges support of Project HIVE 101187384 funded by the European Union. EGB acknowledges partial support for this project from the European Union’s Horizon Europe Research and Innovation programme under grant agreement No. 101134200 “FORSAID: Forest surveillance with artificial intelligence and digital technologies”. Views and opinions expressed are however those of the author(s) only and do not necessarily reflect those of the European Union or the European Research Executive Agency. Neither the European Union nor the granting authority can be held responsible for them.

## Data availability statement

All data supporting the results of this study are included within the article and its Supporting Information files.

